# A Uropathogenic *E. coli* UTI89 model of prostatic inflammation and collagen accumulation for use in studying aberrant collagen production in the prostate

**DOI:** 10.1101/2020.08.10.238196

**Authors:** Hannah Ruetten, Jaskiran Sandhu, Brett Mueller, Peiqing Wang, Helen L. Zhang, Kyle A. Wegner, Mark Cadena, Simran Sandhu, Lisa Abler, Jonathan Zhu, Chelsea A. O’Driscoll, Britta Chelgren, Zunyi Wang, Tian Shen, Jonathan Barasch, Dale E. Bjorling, Chad M. Vezina

## Abstract

Bacterial infection is one known etiology of prostatic inflammation. Prostatic inflammation is associated with prostatic collagen accumulation and both are linked to progressive lower urinary tract symptoms in men. We characterized a model of prostatic inflammation utilizing transurethral instillations of *E. coli* UTI89 in C57BL/6J male mice with the goal of determining the optimal instillation conditions, understanding the impact of instillation conditions on urinary physiology, and identifying ideal prostatic lobes and collagen 1a1 prostatic cell types for further analysis. The smallest instillation volume tested (50 µL) distributes exclusively to bladder, 100 and 200 µL volumes distributes to bladder and prostate, and a 500 µL volume distributes to bladder, prostate and ureter. A threshold optical density (OD) of 0.4 *E. coli* UTI89 in the instillation fluid is necessary for significant (p < 0.05) prostate colonization. *E. coli* UTI89 infection results in a low frequency, high volume spontaneous voiding pattern. This phenotype is due to exposure to *E. coli* UTI89, not catheterization alone, and is minimally altered by a 50 µL increase in instillation volume and doubling of *E. coli* concentration. Prostate inflammation is isolated to the dorsal prostate and is accompanied by increased collagen density. This is partnered with increased density of PTPRC+, ProCOL1A1+ co-positive cells and decreased density of ACTA2+, ProCOL1A1+ co-positive cells. Overall, we determined that this model is effective in altering urinary phenotype and producing prostatic inflammation and collagen accumulation in mice.

## INTRODUCTION

Most men of advancing age experience lower urinary tract symptoms (LUTS), which are characterized by frequent voiding, especially at night (nocturia), weak urine stream, and a feeling of incomplete bladder emptying (20, 35). LUTS reduce quality of life in a significant population of men (10, 16, 38, 53). Men who have LUTS can progress to experiencing severe lower urinary tract dysfunction including urinary retention, bladder wall remodeling, recurrent urinary tract infections, bladder calculi, and renal impairment (20, 22).

Current evidence suggests LUTS derive from a multifactorial process, with prostatic inflammation serving as a key driver (7). Histologic inflammation, especially lymphocytic (chronic) inflammation, is common in men with LUTS and strongly associates with LUTS severity (33). Prostatic inflammation also associates with prostatic collagen abundance, which itself associates with urinary dysfunction (3, 9, 14, 15, 26, 30, 47).

Mice have emerged as an important resource for modeling prostatic inflammation and for identifying molecular signaling pathways that contribute to pathology and urinary voiding physiology (4, 25, 44). Rodent prostate inflammation has been driven by a genetic approach (2), by initiating autoimmunity against the prostate (17, 40), and by introducing noxious stimuli into the prostate (13, 23, 29). Bacterial infection is also used to initiate prostate inflammation, especially in genetically modified mice where it can facilitate a mechanistic understanding of inflammation, pain, collagen accumulation, and voiding dysfunction. Inflammation of the mouse prostate in a transurethral bacterial infection model recapitulates a natural cause of prostatic inflammation and urinary dysfunction in men and dogs (1, 32). The overarching goal of this study is to address some of the remaining questions about this model system, including which bacterial instillation methods are most efficient and reproducible in driving prostate inflammation in mice, if urinary function is altered acutely (less than seven days post bacterial instillation) in C57Bl/6J mice (the most common background strain for commercially available transgenic mice), and which prostatic cells produce collagen in the acute phase of bacterial inflammation. Answering these three questions will facilitate our future goals of determining targetable collagen production pathways and conducting preclinical testing of novel therapeutics.

Bacterial inflammation of the mouse prostate is achieved by catheterizing anesthetized male mice and instilling a bacterial solution into the male urethra. The first goal of this study is to determine how instillation volume and bacterial concentration influence instillation fluid distribution across the lower urinary tract and infection severity within the mouse prostate. We found that large (200 µL+) instillation fluid volumes distribute more widely across the lower urinary tract than a smaller volume (50 µL), which distributes only to the bladder. We found that a threshold concentration of *E. coli* UTI89 in instillation fluid (OD 0.4) is needed to achieve a significant *E. coli* load in the prostate. Further increases, beyond the threshold concentration of *E. coli* in instillation fluid, do not significantly increase the bacterial load in the prostate. We also found that free catch urine, captured and cultured within 24 hours of bacterial instillation, can be used to predict which mice will have histological prostatic inflammation at seven days post installation.

The second goal of this study is to determine the impact of acute *E. coli* UTI89 infection on urinary physiology using the spontaneous void spot assay (VSA) and anesthetized cystometry. Whether catheterization alone induced injury and altered VSA and anesthetized cystometry parameters was previously unknown. We found that urinary catheter placement does not by itself change male mouse urinary function, nor does urethral instillation of sterile saline (up to 100 µL). *E. coli* UTI89 infection results in a low frequency, high volume spontaneous voiding phenotype, which is uniquely observed when monitoring spontaneous voiding and not when monitoring urinary voiding of anesthetized mice by cystometry. Our results are consistent with an *E. coli*-mediated change in mouse behavior and not *E. coli* mediated changes to the lower urinary tract physiology. In addition, there are no differences in voiding behaviors between mice colonized with *E. coli* (positive urine culture) and mice that were uncolonized (negative urine culture) at seven days post instillation, suggesting that voiding behavior changes evoked by transurethral instillation of bacteria are caused by bacterial exposure and not bacterial colonization.

The third goal of this study is to establish anatomical relationships between bacterial colonization, inflammation, and collagen density in the mouse prostate and determine which prostatic cell types produce collagen in inflamed regions. Previous *E. coli* driven mouse models of prostatic inflammation have documented prostatic collagen accumulation (4, 54). No previous study has examined if the *E. coli* UTI89 strain also causes prostatic collagen accumulation and if infection, inflammation, and collagen accumulation are colocalized. A clear relationship has been established between collagen content, prostate stiffness, voiding symptom severity, and resistance to medical therapy in men (26, 27). However, no previous study has determined which cell types are responsible for driving inflammation-mediated prostatic collagen *in vivo*. We evaluated prostate tissue seven days after transurethral delivery of *E. coli* UTI89. The inflammation is confined to the dorsal prostate and is heterogeneous. Collagen is denser in *E. coli* UTI89 infected mice, particularly in regions with dense prostatic stromal cellularity. We also found that the majority of prostatic procollagen 1a1 immunopositive cells within inflammatory lesions are also protein tyrosine phosphatases, receptor type C (also known as CD45) immunopositive and smooth muscle actin immunonegative, suggesting circulating bone marrow derived as a potential origin of prostatic collagen producing cells.

## MATERIALS AND METHODS

### Mice

All experiments were conducted under an approved protocol from the University of Wisconsin Animal Care and Use Committee and in accordance with the National Institutes of Health Guide for the Care and Use of Laboratory Animals. Mice were housed in Udel® Polysulfone microisolator cages on racks or in Innocage® disposable mouse cages on an Innorack®; room lighting was maintained on 12 h light and dark cycles; room temperature was typically 20.5 ± 5°C; humidity was 30–70%. Mice were fed 8604 Teklad Rodent Diet (Harlan Laboratories, Madison WI) and feed and water were available ad libitum. Cages contained corn cob bedding.

To determine the anatomical regions into which instillation fluid distributes (Figure 1), we tested multiple instillation volumes using mice of multiple ages and genetic backgrounds (including retired breeders, old stock animals, or genotype inappropriate animals from other experiments) to follow the 3Rs of humane animal research and reduce animals utilized for this study. Each instillation volume included one C57BL/6J mouse to ensure no variations due to background strain. All mice were sexually mature adult males and included wildtype C57BL/6J #000664, wildtype FVB/NJ #001800, BALB/c-Tg(S100a4-cre)1Egn/YunkJ #012641, Wnt10b-G2aCE #028116, B6.Cg-Tg(Gt(ROSA)26Sor-EGFP)I1Able/J (Rosa-GFP) #007897, B6.Cg-Gt(ROSA)26Sortm14(CAG-tdTomato)Hze/J #007914, 129S/Sv-Ccn2tm2Mae/J #030767, and Tet-ON IL-1b (2).

**Figure 1.**
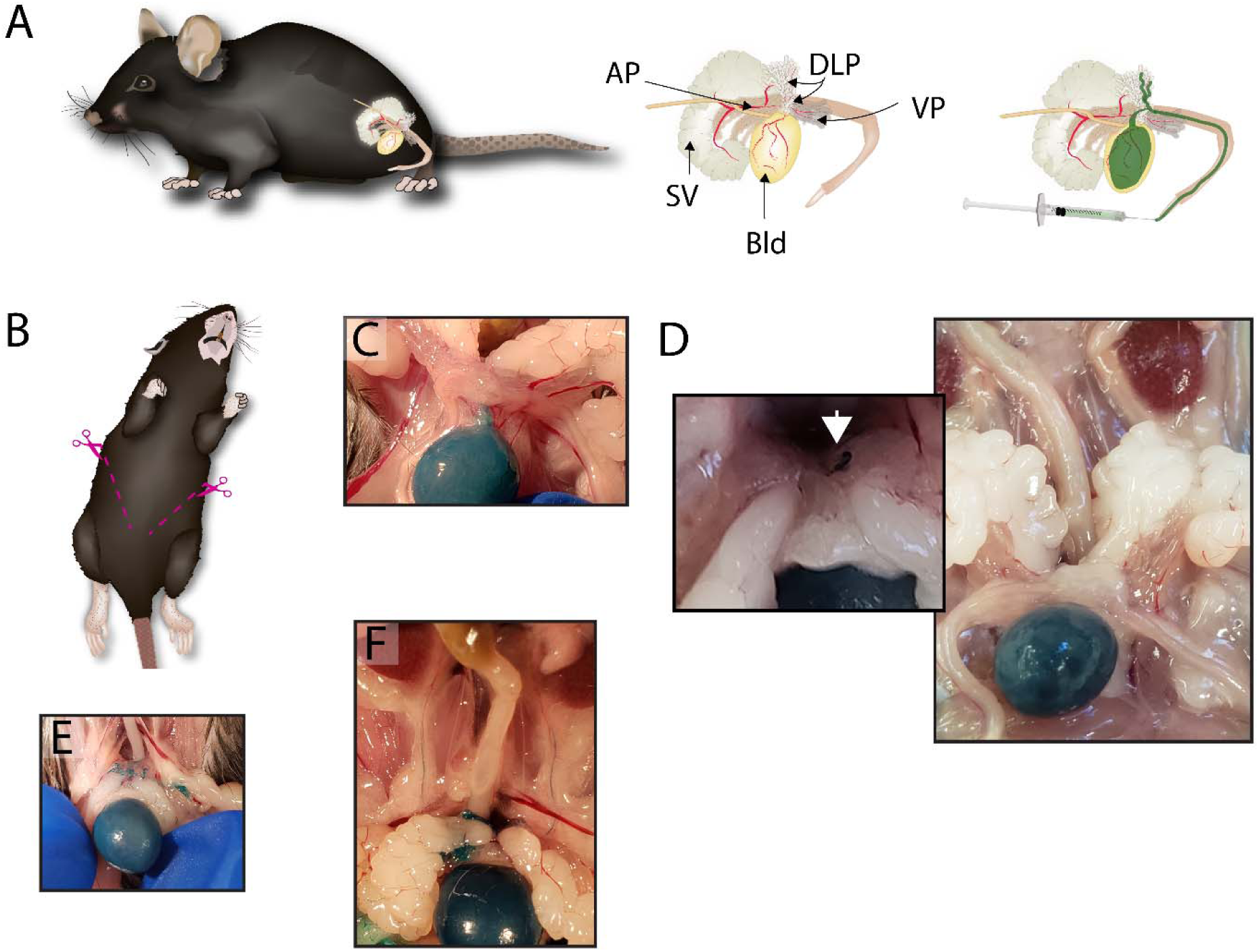
The volume of transurethral instillation fluid determines the anatomical distribution of instilled fluid across the male mouse lower urinary tract. (A) Mouse with lower urinary tract schematic and labeled lower urinary tract anatomical components, including anterior-AP, ventral-VP, dorsolateral-DLP, bladder-Bld, and seminal vesicle-SV. Mice were euthanized, a transurethral catheter (2.5 cm, P10) was placed into mice of various genetic backgrounds and strains (see Materials and Methods) and green Davidson® Tissue Dye was instilled. A loose suture was placed around the tip of the penis, the catheter removed, and suture tightened to prevent dye leakage. (B) Two cuts extending from pubis to lateral ribs were used to open the abdomen and visualize the urinary tract. Ureters, bladder, seminal vesicles, prostatic lobes, and ductus deferens were visually inspected for green dye. (C) Instillation of 50 µL of fluid distributed dye to the bladder. (D) Instillation of 100 µL of fluid distributed dye to the bladder and dorsal prostate. (E) Instillation of 200 µL of fluid distributed dye to the bladder and multiple prostate lobes. (F) Instillation of 500 µL distributed dye to the bladder all prostate lobes, seminal vesicle, and ureters.

C57BL/6J mice (Stock #000664) were used to characterize urinary physiology, stromal cell infiltration, collagen accumulation, and collagen producing cell types (Figures 2, 4-8, and Tables 1-4). C57BL/6J mice were purchased from Jackson Laboratories (Bar Habor, ME) and transurethrally catheterized at six to eight weeks of age.

**Figure 2.**
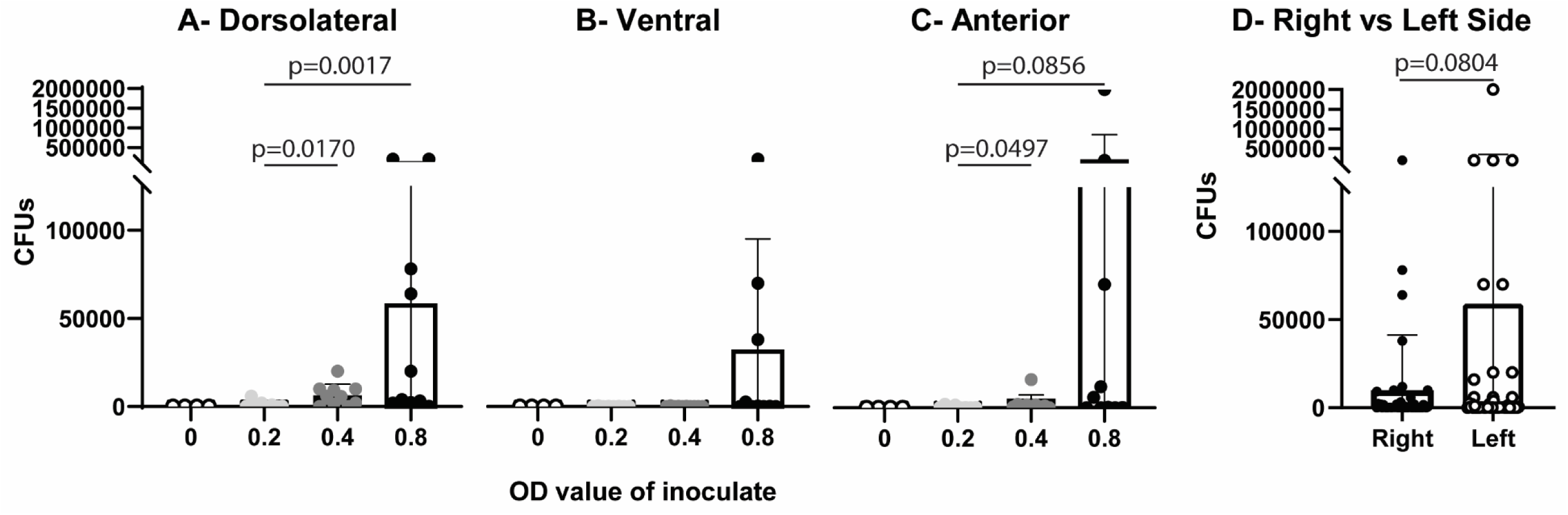
Concentration of *E. coli* in instillation fluid determines acute prostatic CFU load. C57BL/6J male mice were instilled with sterile PBS (optical density-OD 0), or PBS containing graded concentrations of *E. coli* (OD 0.2, 0.4, or 0.8). Mice were euthanized within 1 minute of instillation. Prostate lobes were collected, homogenized, and plated in serial dilution to determine CFUs per mL tissue homogenate. Graphs show the mean ± SD. The mouse prostate contains bilaterally symmetrical prostate lobes and for the purposes of this figure half of one lobe (hemi prostate) was used as the statistical unit. Results are representative of 4-10 hemi prostate lobes from 2-5 mice per group and two hemi prostate lobes per mouse. For A-C, the Shapiro-Wilk test was used to test for normality and transformation was applied to normalize data. Bartlett’s test was used to test for homogeneity of variance. Welsh’s ANOVA was applied when variance was unequal followed by Dunnett’s T3 multiple comparisons test. When variance was equal, comparisons between groups were made using ordinary one-way ANOVA followed by Tukey’s multiple comparisons test. For D, Wilcoxon matched-pairs signed rank test was performed to compare right to left in each lobe (dorsolateral, ventral, anterior) in each mouse instilled with inoculate (OD 0.2-0.8). A p < 0.05 was considered statistically significant. P < 0.10 are shown.

**Figure 3.**
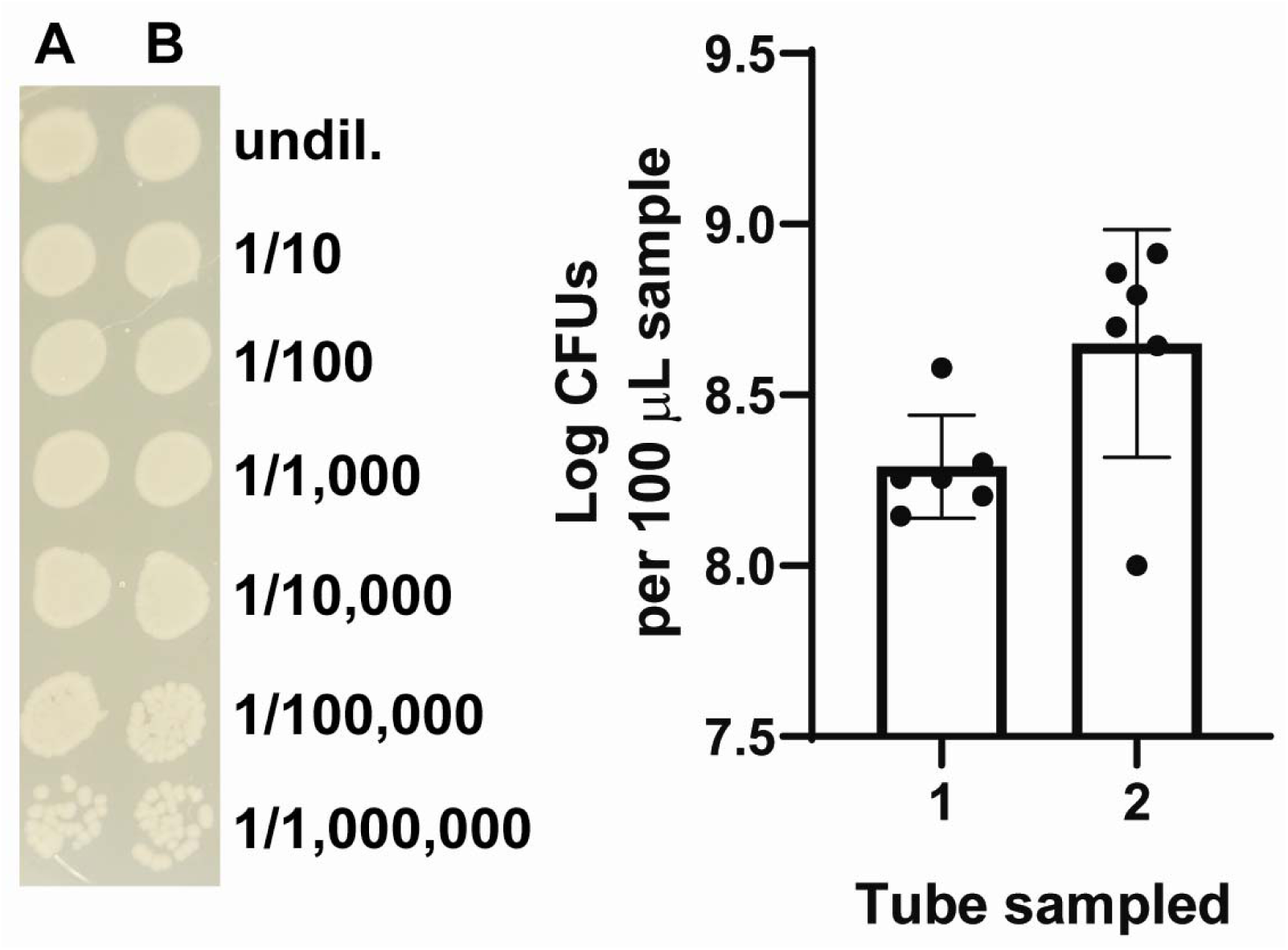
100 µL of *E. coli* UTI89 (OD 0.80) contains 1-8.2 x10^8^ CFUs of *E. coli*. To determine the range of CFUs for *E. coli* UTI89 culture and transurethral instillation, and to establish biologically significant cutoffs for CFUs in tissue or urine, we measured the concentration of twelve (six per tube) instillation aliquots (A-L) originating from two separate culture tubes (Tube 1 and Tube 2). Culture tubes of inoculation solution were prepared at an OD of 0.80 and were prepared by the same person on the same day and both colonies were taken from a single streaked plate. An image of the culture plate of serial dilutes from two aliquots are depicted to the left of the graph. Graph mean ± SD. Shapiro-Wilk test was used to test for normality and transformation was applied to normalize data. Data could not be normalized using transformation, so the Mann Whitney test was applied to compare tubes. A p < 0.05 was considered statistically significant.

**Figure 4.**
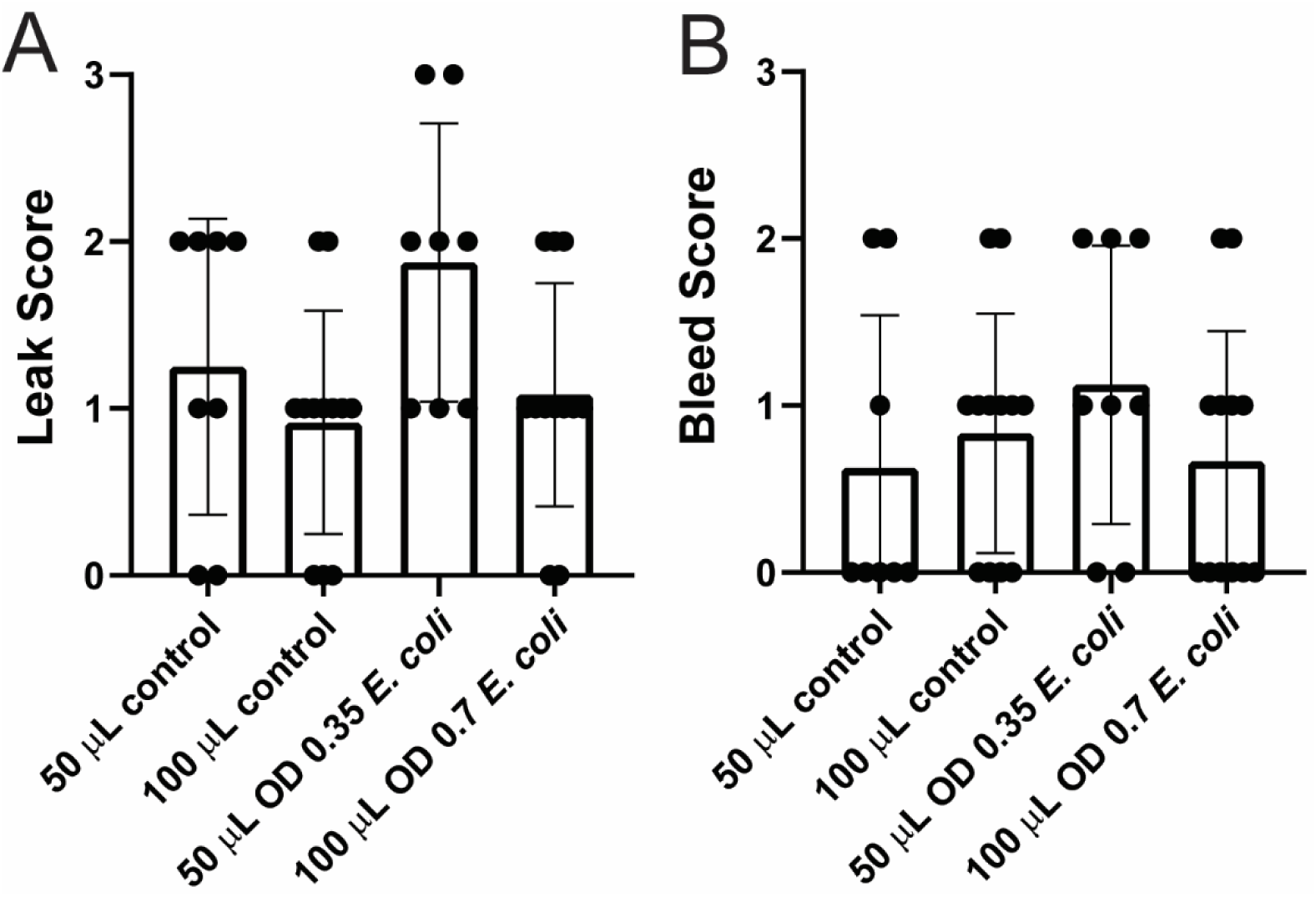
Larger inoculation volumes and *E. coli* UTI89 concentrations do not change the incidence of bleeding or leaking associated with catheter placement. A transurethral catheter was placed in C57BL/6J male mice to deliver either 50 µL of sterile PBS containing *E. coli* UTI89 (OD 0.35) or 100 µL of sterile PBS containing *E. coli* UTI89 (OD 0.7). The methods were compared to determine the influence of inoculation method on the incidence of catheter associated bleeding and leaking. We used a scale of 0-3 to score leaking of instillation fluid from the catheter and urethral bleeding associated with catheterization. For leaking: 0-absent, 1 <10%, 2 <50%, and 3≥ 50% of instilled volume. For bleeding: 0-absent, 1-minimal, but no continuous bleeding, 2-minimal continued bleeding, 3-continued bleed that requires holding pressure. Graphs are mean ± SD and representative of n = 8-12 per group. Groups were compared using a Kruskal-Wallis test. A p < 0.05 was considered statistically significant.

**Table 1.** Spontaneous void behavior 1d post urethral instillation with sterile PBS (control) or PBS containing *E. coli* UTI89.

| Void Spot Assay | Values are: Average (SD) |  |  |  |  |  |
| --- | --- | --- | --- | --- | --- | --- |
| | 50 μL<br>Sterile PBS<br>Control<br>(n = 8) | 100 μL<br>Sterile PBS<br>Control<br>(n = 12) | All<br>Sterile PBS<br>Control<br>(n = 22) | 50 μL<br>PBS w/ OD 0.35<br>*E. coli* UTI89<br>(n = 8) | 100 μL<br>PBS w/ OD 0.7<br>*E. coli* UTI89<br>(n = 12) | All<br>PBS w/ *E. coli*<br>UTI89<br>(n=22) |
| Count | 15.25 (10.87) | 21.92 (36.15) | 19.77 (27.16) | 6.125 (4.970) | 13.17 (13.60) | 13.23 (14.27) |
| Total Area (cm<sup>2</sup>) | 18.32 (11.88) | 16.54 (12.64) | 16.67 (11.80) | 13.45 (16.49) | 12.45 (6.532) | 15.03 (12.91) |
| Percent area in<br>center | 7.907 (21.01) | 3.434 (5.772) | 7.287 (16.90) | 13.61 (31.72) | 14.81 (29.92) | 13.73 (28.50) |
| Percent area in<br>corners | 60.46 (43.76) | 63.27 (28.87) | 59.95 (34.31) | 37.76 (40.05) | 41.37 (40.54) | 39.22 (37.53) |
| Spots of a certain size (cm<sup>2</sup>) |  |  |  |  |  |  |
| 0 - 0.1 | 12.75 (9.867) | 17.17 (29.69) | 15.68 (22.36) | 4.625 (4.069) | 11.67 (13.21) | 11.41 (13.32) |
| 0.1 - .25 | 0.6250 (1.061) | 2.333 (5.990) | 1.591 (4.469) | 0.3750 (0.5175) | 0.08333 (0.2887) | 0.3636 (0.6580) |
| 0.25 - .5 | 0.5000 (0.5345) | 0.5000 (0.9045) | 0.7727 (1.572) | 0.1250 (0.3536) | 0.08333 (0.2887) | 0.1818 (0.3948) |
| 0.5 - 1 | 0.2500 (0.4629) | 0.5833 (0.9962) | 0.5000 (0.8591) | 0.1250 (0.3536) | 0 (0) | 0.09091 (0.2942) |
| 1 - 2 | 0 (0) | 0.08333 (0.2887) | 0.04545 (0.2132) | 0.1250 (0.3536) | 0 (0) | 0.04545 (0.2132) |
| 2 - 3 | 0 (0) | 0.08333 (0.2887) | 0.04545 (0.2132) | 0 (0) | 0.08333 (0.2887) | 0.04545 (0.2132) |
| 3 - 4 | 0 (0) | 0 (0) | 0.04545 (0.2132) | 0 (0) | 0.1667 (0.5774) | 0.09091 (0.4264) |
| 4+ | 1.125 (0.6409) | 1.167 (0.9374) | 1.091 (0.8112) | 0.7500 (0.8864) | 1.083 (0.5149) | 1.000 (0.6901) |

**Table 2.** Spontaneous void behavior 3d post urethral instillation with sterile PBS (control) or PBS containing *E. coli* UTI89.

| Void Spot Assay | Values are: Average (SD) |  |  |  |  |  |
| --- | --- | --- | --- | --- | --- | --- |
| | 50 μL<br>Sterile PBS<br>Control<br>(n = 8) | 100 μL<br>Sterile PBS<br>Control<br>(n = 11) | All<br>Sterile PBS<br>Control<br>(n = 21) | 50 μL<br>PBS w/ OD 0.35<br>*E. coli* UTI89<br>(n = 8) | 100 μL<br>PBS w/ OD 0.7<br>*E. coli* UTI89<br>(n = 11) | All<br>PBS w/ *E. coli*<br>UTI89<br>(n =21) |
| Count | 136.6 (221.6) | 20.55 (17.03) | 64.19 (144.0) | 41.63 (61.91) | 9.455 (13.37) | 22.43 (40.94) |
| Total Area (cm<sup>2</sup>) | 21.33 (6.374) | 23.40 (14.08) | 21.44 (11.63) | 26.13 (15.49) | 21.75 (13.83) | 25.60 (15.27) |
| Percent area in<br>center | 6.374 (11.03) | 9.612 (15.64) | 8.674 (13.61) | 1.168 (1.619) | 17.77 (31.76) | 9.996 (23.98) |
| Percent area in<br>corners | 61.36 (28.20) | 51.88 (29.38) | 57.20 (28.08) | 46.11 (34.84) | 63.67 (22.24) | 56.28 (26.72) |
| Spots of a certain size (cm<sup>2</sup>) |  |  |  |  |  |  |
| 0 - 0.1 | 116.1 (197.0) | 16.18 (15.4) | 53.71 (127.4) | 34.13 (48.85) | 6.909 (12.45) | 17.95 (32.98) |
| 0.1 - .25 | 14.38 (30.18) | 1.182 (1.662) | 6.238 (19.05) | 3.750 (8.294) | 0.1818 (0.4045) | 1.619 (5.210) |
| 0.25 - .5 | 3.75 (5.825) | 0.5455 (0.6876) | 1.905 (3.807) | 1.375 (2.722) | 0.3636 (0.6742) | 0.7143 (1.765) |
| 0.5 - 1 | 1.250 (2.375) | 0.4545 (0.6876) | 0.7143 (1.554) | 0.3750 (1.061) | 0.2727 (0.4671) | 0.3333 (0.7303) |
| 1 - 2 | 0.2500 (0.7071) | 0.1818 (0.4045) | 0.1905 (0.5118) | 0.2500 (0.4629) | 0.1818 (0.4045) | 0.1905 (0.4024) |
| 2 - 3 | 0 (0) | 0.1818 (0.4045) | 0.09524 (0.3008) | 0.1250 (0.3536) | 0.1818 (0.4045) | 0.1429 (0.3586) |
| 3 - 4 | 0 (0) | 0 (0) | 0.000 (0.000) | 0.1250 (0.3536) | 0.09091 (0.3015) | 0.09524 (0.3008) |
| 4+ | 0.8750 (0.6409) | 1.818 (1.250) | 1.333 (1.111) | 1.500 (0.7559) | 1.273 (0.7862) | 1.381 (0.7400) |

**Table 3.** Spontaneous void behavior 5d post urethral instillation with sterile PBS (control) or PBS containing *E. coli* UTI89.

| Void Spot Assay | Values are: Average (SD) |  |  |  |  |  |
| --- | --- | --- | --- | --- | --- | --- |
| | 50 μL<br>Sterile PBS<br>Control<br>(n = 8) | 100 μL<br>Sterile PBS<br>Control<br>(n = 11) | All<br>Sterile PBS<br>Control<br>(n = 21) | 50 μL<br>PBS w/ OD 0.35<br>*E. coli* UTI89<br>(n = 8) | 100 μL<br>PBS w/ OD 0.7<br>*E. coli* UTI89<br>(n = 11) | All<br>PBS w/ *E. coli*<br>UTI89<br>(n = 21) |
| Count | 41.50 (40.10) | 19.50 (24.95) | 27.18 (31.55) | 38.13 (22.64) | 12.55 (20.87) | 23.29 (23.43) |
| Total Area (cm<sup>2</sup>) | 23.76 (7.168) | 14.11 (9.768) | 17.59 (10.32) | 25.20 (12.69) | 28.26 (19.41) | 26.78 (16.04) |
| Percent area in center | 8.175 (16.53) | 1.611 (3.119) | 7.919 (20.87) | 3.491 (4.650) | 4.979 (8.877) | 3.961 (6.942) |
| Percent area in corners | 65.38 (27.77) | 75.18 (29.03) | 67.81 (30.83) | 34.65 (27.82) | 48.81 (36.18) | 46.05 (32.77) |
| Spots of a certain size (cm<sup>2</sup>) |  |  |  |  |  |  |
| 0 - 0.1 | 30.75 (24.45) | 16.92 (24.04) | 21.68 (23.69) | 33.50 (19.49) | 9.909 (20.94) | 19.81 (21.94) |
| 0.1 - .25 | 5.750 (12.68) | 0.9167 (1.311) | 2.636 (7.768) | 1.500 (1.852) | 0.1818 (0.4045) | 0.8095 (1.401) |
| 0.25 - .5 | 1.625 (3.021) | 0.2500 (0.8660) | 0.7727 (1.974) | 1.250 (1.753) | 0.1818 (0.4045) | 0.6190 (1.203) |
| 0.5 - 1 | 1.125 (1.727) | 0.1667 (0.3892) | 0.5455 (1.143) | 0.2500 (0.4629) | 0.3636 (0.5045) | 0.2857 (0.4629) |
| 1 - 2 | 0.1250 (0.3536) | 0 (0) | 0.04545 (0.2132) | 0 (0) | 0 (0) | 0.000 (0.000) |
| 2 - 3 | 0 (0) | 0.08333 (0.2887) | 0.04545 (0.2132) | 0 (0) | 0 (0) | 0.000 (0.000) |
| 3 - 4 | 0.1250 (0.3536) | 0.08333 (0.2887) | 0.09091 (0.2942) | 0 (0) | 0 (0) | 0.000 (0.000) |
| 4+ | 2.000 (1.195) | 1.083 (0.6686) | 1.364 (1.002) | 1.625 (0.9161) | 1.909 (1.044) | 1.762 (0.9437) |

**Table 4.** Spontaneous void behavior 7d post urethral instillation with sterile PBS (control) or PBS containing *E. coli* UTI89.

| Void Spot Assay | Values are: Average (SD) |  |  |  |  |  |
| --- | --- | --- | --- | --- | --- | --- |
| | 50 μL<br>Sterile PBS<br>Control<br>(n = 8) | 100 μL<br>Sterile PBS<br>Control<br>(n = 12) | All<br>Sterile PBS<br>Control<br>(n = 21) | 50 μL<br>PBS w/ OD 0.35<br>*E. coli* UTI89<br>(n = 7) | 100 μL<br>PBS w/ OD 0.7<br>*E. coli* UTI89<br>(n = 11) | All<br>PBS w/ *E. coli*<br>UTI89<br>(n = 20) |
| Count | 19.75 (17.65) | 46.67 (78.86) | 40.81 (64.88) | 37.00 (21.39) | 7.273 (10.43) | 19.50 (20.53) |
| Total Area (cm<sup>2</sup>) | 11.26 (16.11) | 17.54 (10.25) | 15.94 (13.25) | 26.14 (14.10) | 29.38 (13.37) | 28.47 (13.25) |
| Percent area in<br>center | 1.936 (2.531) | 5.121 (8.867) | 4.554 (7.643) | 8.606 (15.36) | 7.314 (9.500) | 7.159 (11.25) |
| Percent area in<br>corners | 44.74 (38.97) | 63.44 (31.63) | 55.72 (34.16) | 54.14 (31.55) | 55.83 (29.51) | 58.00 (29.16) |
| Spots of a certain size (cm<sup>2</sup>) |  |  |  |  |  |  |
| 0 - 0.1 | 17.75 (15.79) | 37.08 (64.06) | 31.81 (50.60) | 31.57 (18.12) | 3.909 (9.016) | 15.20 (18.28) |
| 0.1 - .25 | 0.6250 (1.061) | 5.583 (13.83) | 5.143 (12.71) | 2.000 (3.162) | 0.3636 (0.9244) | 1.050 (2.064) |
| 0.25 - .5 | 0.1250 (0.3536) | 1.333 (2.229) | 1.381 (3.008) | 1.286 (1.254) | 0 (0) | 0.5000 (0.9459) |
| 0.5 - 1 | 0.2500 (0.7071) | 0.4167 (0.6686) | 0.6667 (1.592) | 0.4286 (0.5345) | 0 (0) | 0.1500 (0.3663) |
| 1 - 2 | 0 (0) | 0.3333 (0.7785) | 0.2857 (0.7171) | 0 (0) | 0.09091 (0.3015) | 0.1000 (0.3078) |
| 2 - 3 | 0.1250 (0.3536) | 0.4167 (0.7930) | 0.2857 (0.6437) | 0 (0) | 0 (0) | 0.05000 (0.2236) |
| 3 - 4 | 0.1250 (0.3536) | 0.8333 (0.2887) | 0.09524 (0.3008) | 0 (0) | 0.1818 (0.4045) | 0.1500 (0.3663) |
| 4+ | 0.7500 (1.165) | 1.417 (0.9962) | 1.143 (1.062) | 1.714 (0.7559) | 2.727 (0.7862) | 2.300 (0.9234) |

### E. coli

Experiments involved *E. coli* UTI89 (31), a strain of uropathogenic *Escherichia coli* (UPEC) recovered from a human patient with cystitis (24) and transformed with a pCOMGFP plasmid (48) to confer green fluorescent protein expression and kanamycin resistance. Prior to inoculation, *E. coli* UTI89 were grown as a static culture for 18 hours in antibiotic free Luria-Bertani broth at 37°C. The optical density (OD) was determined prior to mouse inoculation. The culture was centrifugated for 15 minutes at 1157 rcf and the resulting pellet was resuspended into sterile PBS for instillation.

### Transurethral instillation

At six to eight weeks of age, C57BL/6J mice were anesthetized with isoflurane and instilled with sterile PBS (control) or PBS containing *E. coli* UTI89 via transurethral catheter as previously described (25).

### Tissue dye instillation

Mice were euthanized, a transurethral catheter (2.5 cm, PE10) was placed and green Davidson® Tissue Dye was instilled (Figure 1A). A loose suture was placed around the tip of the penis, the catheter removed, and suture tightened to prevent dye leakage. Two cuts extending from pubis to lateral ribs were used to open the abdomen and visualize the urinary tract (Figure 1B). Ureters, bladder, seminal vesicles, prostatic lobes, and ductus deferens were visually inspected for green dye. The instillation fluid was considered to have distributed to a lower urinary tract compartment if staining was visibly apparent in any part of that compartment, including a single duct of the prostate lobe.

### Free catch urine culture

Mice were restrained using a two-finger tail grip and placed in a squat posture over a sterile petri dish. Most mice urinated immediately. Those that did not urinate were scruffed with the handlers opposite hand while in the squat posture. If the mouse still did not urinate, the handler rotated the scruffing hand so the mouse was in dorsal recumbency in the palm of the hand. The handler’s thumb and index finger were swept from the lateral abdomen (near kidneys) toward the pubis applying slight pressure to locate and apply pressure to the bladder. The mouse was positioned above the petri dish and the bladder expressed. Collected urine was serial diluted in sterile PBS and plated on Luria-Bertani (LB) agar containing kanamycin (KAN; 50 µg/mL). Culture Colony forming units (CFU) were calculated per mL of urine.

### Prostate tissue culture

Mice were euthanized, place on sterile surgical drape, and drenched in 70% EtOH. Because our goal was to determine the distribution and severity of *E. coli* infection, great care was taken during dissection to reduce the risk of cross contaminating organs or surgical instruments. A sterile scissors was used to open the body cavity and expose the lower urinary tract. Intestines were moved aside with a sterile gloved finger. Needle tipped forceps were used to grasp the bladder and pull up while a second sterile scissors was used to sever the bladder ligament and the vas deferens. The same forceps were used to pass sterile silk suture under the dorsal surface of the urethra at the pelvic inlet. Hand ties were placed to close the urethra. The same scissors utilized to cut the bladder ligament was used to sever the urethra caudal to the suture tie. The strings of the urethra suture were used to gently exteriorize the bladder and attached seminal vesicle and prostate lobes from the abdomen and place in a sterile petri dish containing sterile PBS. The ureters were severed due to tension during removal. The same forceps used to grasp the bladder earlier was used to adjust the position of the tissue in the dish. A new clean needle tipped forceps was used to remove the lobes by grasping the urethral attachment and pulling. The left lobes were removed first, ventral lobe, then dorsolateral, and finally anterior, then right lobes in the same order. Each lobe was placed in a sterile 1.5 mL tube with 100 µL of sterile PBS and hand ground with a sterile pestle. Supernatant was removed and placed in a 96-well plate for serial dilution (10-fold dilution series) and 5 µL was plated per dilution on LB agar containing KAN (50 µg/mL). Scissors and forceps were sterilized between mice using a bead sterilizer and a new sterile surgical drape, sterile suture, sterile petri dish, and sterile PBS were used for each mouse. CFUs were calculated per mL of supernatant.

### Urinary Function Testing

Void spot assay (VSA) and cystometry were performed and analyzed as previously described (42). We followed the recommended guidelines of reporting VSA data (19, 21, 51). VSA was performed in the vivarium where mice were housed. Whatman grade 540 (Fisher Scientific no. 057163-W) filter papers (27 × 16 cm) were placed in the bottom of Udel^®^ Polysulfone microisolator cages. Mice were placed in the cage (singly housed) with food *ad libitum* but no water for four hours starting from 8-11 AM GMT. Filter papers were dried and imaged with an Autochemi AC1 Darkroom ultraviolet imaging cabinet (UVP, Upland, CA) equipped with an Auto Chemi Zoom lens 2UV and an epi-illuminator. Image capture settings were adjusted using UVP VisonWorksLS image acquisition software. Images were captured using an Ethidium Bromide filter set (570-640 nm) and 365 nm epi-illumination. Void Whizzard was downloaded from http://imagej.net/Void_Whizzard and run according to the user guide (51). Analyzed parameters included: Total Spot Count, Total Void Area (cm^2^), % area in center of paper, % area in corners of paper, and mass distribution of spots (0-0.1, 0.1-0.25, 0.25-0.5, 0.5-1, 1-2, 2-3, 3-4, 4+ cm^2^).

Cystometry was performed with minimal alterations to previously published protocols and following best practices (5, 12, 37). Mice were anesthetized with urethane (1.43 g/kg, s.c.). Thirty minutes after urethane dosing, an incision was made in the ventral abdomen to expose the bladder. Bladder length and diameter were measured for volume calculation. A purse-string suture was placed in the bladder dome. Polyethylene cystostomy tubing (PE50, outer diameter 0.965 mm, inner diameter 0.58 mm) was inserted into the bladder through the center of the suture and purse-string secured to hold the tubing in place with 2-3 mm of tubing within the bladder. The abdominal wall and skin were closed separately in a simple interrupted pattern. The exterior tubing was secured to the ventral abdominal skin with two simple interrupted sutures. Mice were placed on a heat pad for one hour after the procedure.

The exposed cystostomy tube was connected to a three-way stopcock, and the other two arms of the stopcock were connected to an infusion pump (Harvard Apparatus, Holliston, MA) and pressure transducer (Memscap AS, Norway). Intravesical pressure was recorded continuously using a PowerLab data collection system (ADI Instruments, Colorado Springs, CO). Room-temperature sterile saline (0.9%) was infused into the bladder at a rate of 0.8 mL per hour.

Mice were placed in lateral recumbency above a force transducer (Model FT03, grass Instruments) attached to a 3D printed urine collection funnel. The force transducer was calibrated with known volumes of saline to create a pressure-volume conversion. The mass of voided urine was recorded continuously using PowerLab.

At least one hour of voiding activity was recorded. Three to five consecutive voids, occurring after stabilization of micturition cycles, were used for analyses. Multiple parameters were measured as previously defined (42) and included: Void Duration, Intervoid Interval, Baseline Pressure, Normalized Threshold Pressure, Normalized Peak Void Pressure, Number of Non-Voiding Contractions, Voided Volume, Compliance, Volume Flow Rate, Mass Based Flow Rate, and Efficiency.

### Tissue preparation

Lower urinary tracts were collected for histology seven- or eight-days post-instillation. Tissues were prepared, fixed, and sectioned as described previously (28, 49, 50). To remove the lower urinary tract, ureters were cut at the entry to the bladder wall, vas deferens cut at the entry to the bladder neck, and urethra was cut immediately dorsal to the pubic symphysis. Tissues were prepared for embedding in one of three ways. For the first method, seminal vesicles, ampullary gland, anterior prostate, and hemi-ventral, dorsal, and lateral prostates were removed. Bladder and urethra were separated at level of the bladder neck with a single transverse cut. Urethras with hemi-ventral, dorsal, and lateral prostates attached were fixed in 4% paraformaldehyde, washed in PBS, and placed in arrays using 1% bactoagar. The array was dehydrated in ethanol, cleared in xylene, and infiltrated with paraffin. Urethras were serially sectioned starting at the bladder neck in the transverse plane (41). For the second method, hemi-dorsal prostate lobes were removed, fixed in 4% paraformaldehyde, washed in PBS, and placed in arrays using 1% bactoagar. The array was dehydrated in ethanol, cleared in xylene, and infiltrated with paraffin. The lobes were sectioned in the plane yielding the greatest surface area of dorsal prostate. For the third method, hemi-seminal vesicle and prostate lobes were removed as well as ampullary gland. Remaining bladder, urethra, and hemi-seminal vesicle and prostate lobes were fixed in 4% paraformaldehyde, washed in PBS, dehydrated in ethanol, cleared in xylene, and infiltrated with paraffin. Sections were cut in the sagittal plane. All sections were five-microns thick and mounted on Superfrost™ Plus Gold Slides (Thermo Fisher Scientific; Waltham, MA).

### Hematoxylin and Eosin (H&E)

H&E staining was performed by the Histology Service in the School of Veterinary Medicine at University of Wisconsin-Madison. Stains were imaged using a BZ-X710 digital microscope (Keyence, Itasca, IL, USA) fitted with at 20x (PlanFluor, NA 0.45) objective. Tiled images were captured using Keyence image acquisition software (Keyence) and stitched to generate an image across the entire section.

### Collagen quantification with Picrosirius Red Staining

Picrosirius red staining (PSR), fluorescent imaging and quantitation was performed as previously described (52). Stained tissue sections were cleared with xylene and mounted with Richard-Allan toluene-based mounting medium. PSR staining was then imaged using a BZ-X710 digital microscope (Keyence, Itasca, IL, USA) fitted with at 20x (PlanFluor, NA 0.45) objective and illuminated by full spectrum light filtered with Texas Red and FICZ filters. Tiled images were captured using Keyence image acquisition software (Keyence) and stitched to generate an image across the entire section.

Specific collagen staining was determined by subtracting the tissue autofluorescence (FICZ channel) from the PSR fluorescence (Texas Red channel) using the “image calculator” function of ImageJ. Additional fluorescence from ductal lumens was removed manually. Total image area was defined as the pixel area of the image. Prostatic stromal area was calculated by tracing exterior boundaries of lobes, and duct lumen, calculating the pixel area and subtracting the area from the total pixel area of the image. Collagen density was defined as the PSR pixel area divided by the stromal tissue area. Analysis was performed on three images per mouse and five to nine mice per group.

### Immunohistochemistry and Cell Counts

Immunohistochemistry (IHC) was conducted on dorsal prostate tissue sections from mice in the sterile PBS control group and the *E. coli* UTI89 infected group. Tissue sections were deparaffinized with xylene and 50%, 75%, and 100% ethanol. Tissues were immersed in citrate buffer pH 6.0 heated in a microwave for epitope decloaking. Tris-buffered saline containing 0.1% Tween 20 and 5% Donkey serum was used as a blocking reagent, and primary and secondary antibodies were diluted in blocking reagent. Antibodies and dilutions are: Procollagen type 1a1 (ProCOL1A1, Developmental Studies Hybridoma Bank, SP1.D8-c (concentrated, 238 µg/mL), 1:1000), Protein tyrosine phosphatase, receptor type C (PTPRC, also known as CD45, Abcam, AB10558, RRID: AB 442810, 1:500), Smooth Muscle Actin Alpha (SMA or ACTA2, Invitrogen, PA5-18292, RRID: AB 10980764, 1:100), Anti-Goat 488 (Jackson ImmunoResearch, 705-545-003, RRID: AB_2340428, 1:250), Anti-Rabbit Cy3 (Jackson ImmunoResearch, 711-165-152, RRID: AB_2307443, 1:250), Anti-Mouse 647 (Jackson ImmunoResearch, 715-605-150, RRID: AB_2340846, 1:250), and DAPI (2-(4-amidinophenyl)- 1H -indole-6-carboxamidine). Sections were imaged using an SP8 Confocal Microscope (Leica, Wetzlar, Germany) fitted with a 20x oil immersion objective (HC PL Apo CS2 NA = 0.75; Leica, Wetzlar, Germany). Samples were excited and detected using the recommended settings for each secondary antibody fluorophore. Images were captured at 1024×1024 resolution using LASX 8 software (Leica, Wetzlar, Germany). Three images of dorsal prostate were captured per mouse and five to nine mice were imaged per group.

Collagen producing cells were defined as ProCOL1A1 immunopositive. ProCOL1A1 is the precursor to collagen 1A1, the most abundant collagen subtype in the extracellular matrix (36). Bone marrow derived cells were defined as PTPRC immunopositive. Myofibroblasts were defined as ACTA2 immunopositive and differentiated from fibroblasts which were defined as ACTA2 immunonegative. DAPI was used to identify cell nuclei. Cells lying within blood vessels, prostate ducts, or prostate ductal epithelium were excluded. All remaining nucleated cells were counted and all cells that were partially or completely stained were considered positive. Cells were manually counted per image using Image J cell counter (43).

### Statistical Analysis

Statistical analyses were performed with Graph Pad Prism 8.0.2. Differences were considered significant at the p < 0.05 level. A Shapiro-Wilk test was used to test for normality and transformation was applied to normalize data. Bartlett’s test was used to test for homogeneity of variance. For group comparisons, Welsh’s ANOVA was applied when variance was unequal followed by Dunnett’s T3 multiple comparisons test. When variance was equal, comparisons between groups were made using ordinary one-way ANOVA followed by Tukey’s multiple comparisons test. A Kruskal-Wallis test was applied when data could not be normalized through transformation. For pair wise comparisons, Student’s t-test was applied when variances were equal and a t-test with Welsh’s correction was applied when variance were unequal between groups. A Mann Whitney test was performed when data could not be normalized through transformation. A Wilcoxon matched-pairs signed rank test was performed for matched pairs. A Fisher’s Exact test was performed to determine if there were associations between infection conditions, catheter induced bleeding and leaking, and biomarkers of prostatic inflammation. Linear regression was performed to correlate stromal cell density with pixel density. For Figure 5 only, a mixed-effects model REML was fit to the data and Geisser-Greenhouse correction applied.

**Figure 5.**
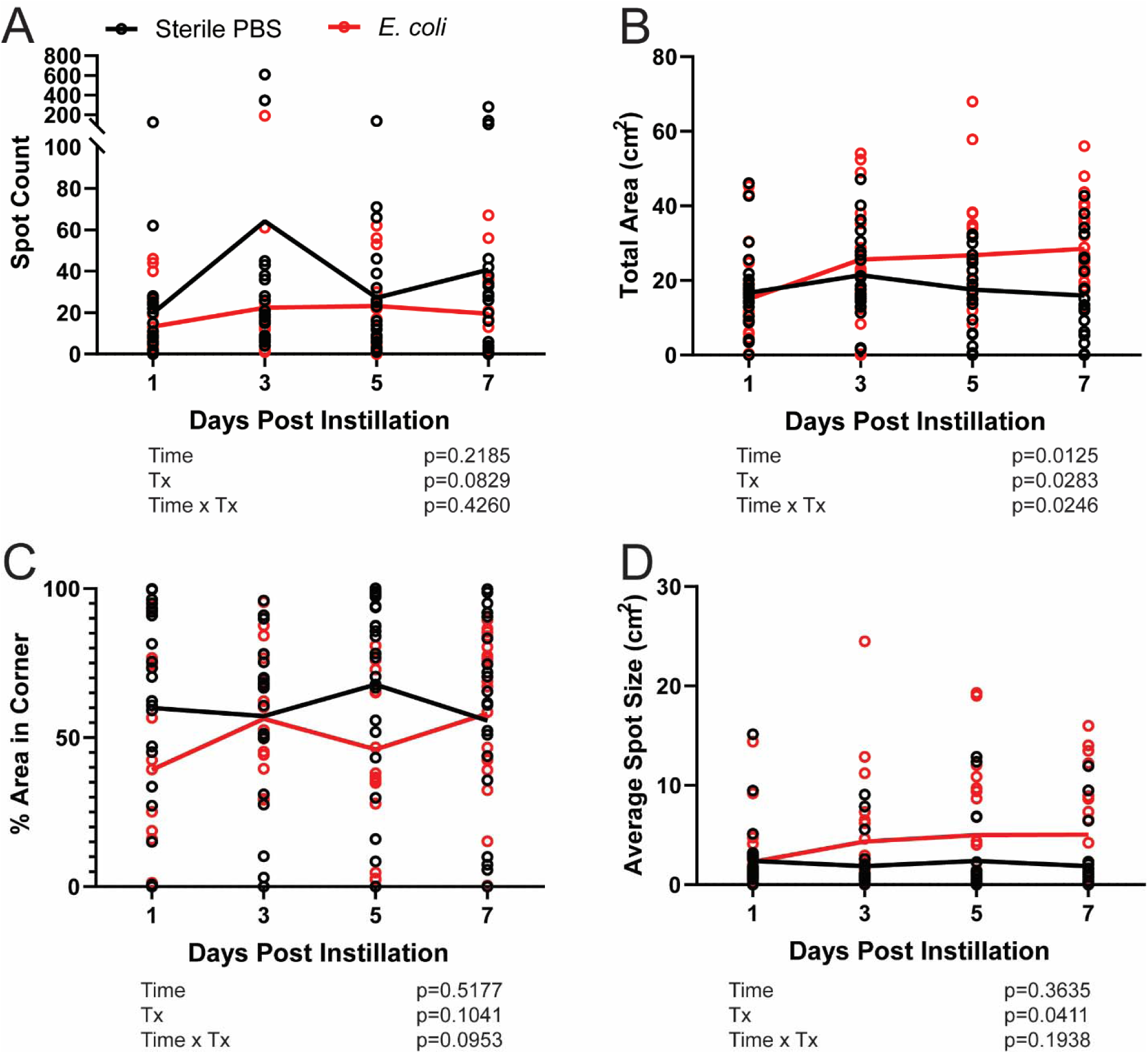
*E. coli* UTI89 infection results in a low frequency, high volume voiding phenotype. All control and *E. coli* UTI89 infected C57BL/6J male mice were grouped, regardless of instillation concentration or volume, to maximize sample size compared spontaneous void spot assays endpoints over time and between *E. coli* UTI89 infected mice and PBS controls. Included are selected endpoints that summarize the detailed void spot assay data included in Tables 1-5. Lines are placed at the mean. A mixed-effects model REML was fit to the data and Geisser-Greenhouse correction applied. Results are representative of n = 20-22 per group. A p < 0.05 was considered statistically significant.

**Table 5.**
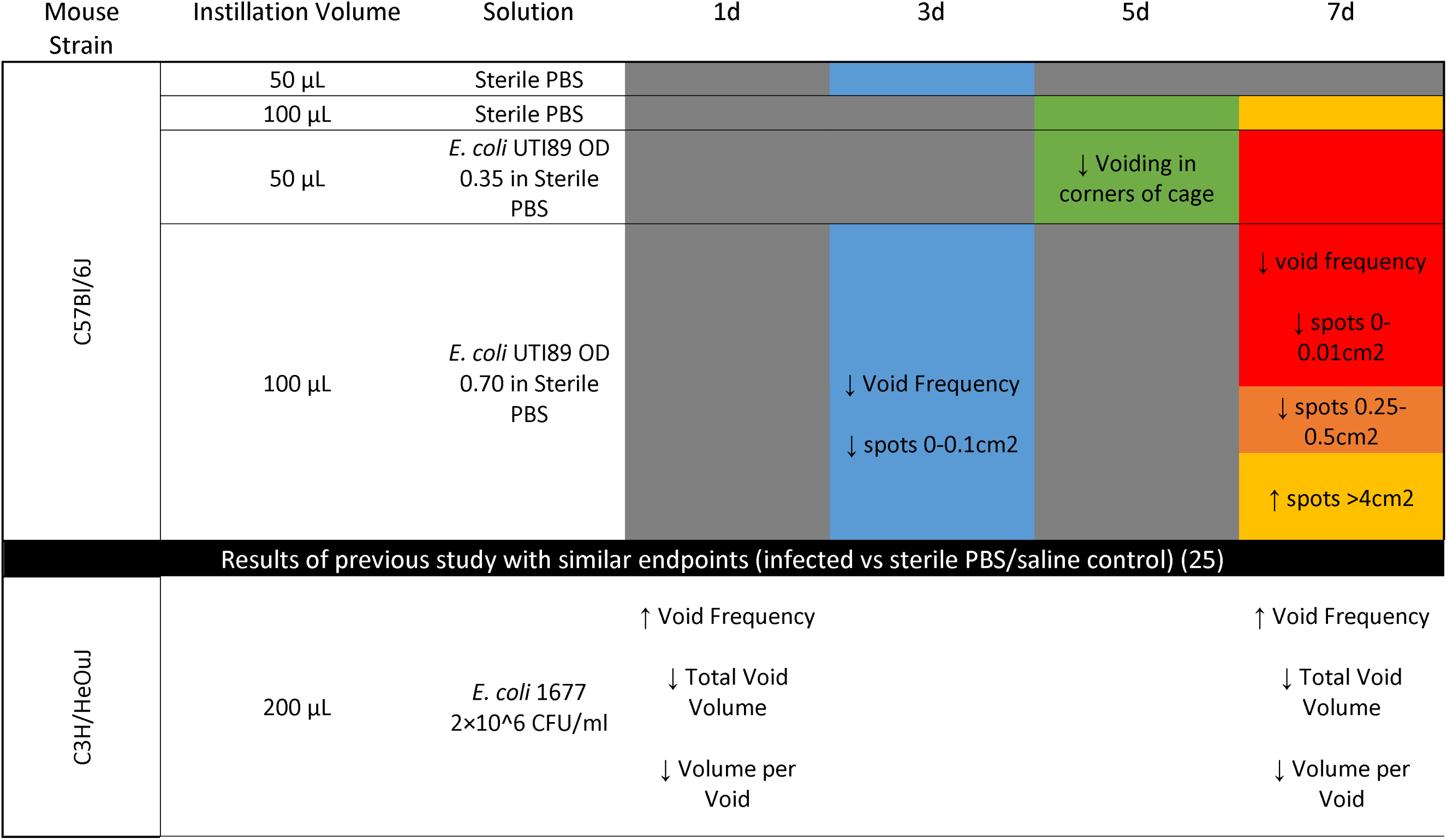

## RESULTS AND DISCUSSION

### The volume of transurethral instillation fluid determines how widely the instilled fluid distributes across the male mouse lower urinary tract

A common goal of urologic researchers using the bacterial infection model is to increase its consistency by identifying factors that drive or enhance prostatic inflammation. Elkahwaji et al. were among the first to deliver bacteria transurethrally to male mice for the purpose of driving prostate inflammation and reported heterogenous and inconsistent infection rates (11). Volumes of instillation fluid in previous studies range from 10-200 µL (6, 39). One reason for the large range of instillation volumes is that it is unclear how bacteria gain access to the prostate. Some believe that large volumes of instilled fluid create hydrostatic pressure and drive bacteria into the prostate. Bacteria could also enter the prostate as reflux in voided urine or actively “crawl” into the prostate through swimming, swarming, or twitching motility.

Here, we test the impact of instilled volume on anatomical distribution of instilled fluid, which may in turn influence the distribution of inflammation. Three mice were instilled with 100 µL of undiluted tissue dye and dye was observed in the bladder (3/3 mice, 100%) and dorsal prostate (3/3 mice, 100%) (Figure 1D). The tissue dye was diluted in PBS to decrease viscosity and delivered via a transurethral catheter into freshly euthanized mice in a volume of 50, 100, 200, and 500 µL (three mice per volume). The 50 µL instillation volume results in the visual appearance of dye in the bladder (3/3 mice, 100%) and minimal infiltration of the dorsolateral prostate (1/3 mice, 33.3%) but not in the anterior (0/3 mice, 0%) or ventral (0/3 mice, 0%) prostate (Figure 1C). The 100 µL instillation volume results in the appearance of dye in the bladder (3/3 mice, 100%) and dorsolateral prostate (3/3 mice, 100%) but not in the anterior (0/3 mice, 0%) or ventral (0/3 mice, 0%) prostate. The 200 µL instillation volume results in the appearance of dye in the bladder (3/3 mice, 100%), dorsolateral (3/3 mice, 100%), anterior prostate (3/3 mice, 100%), and seminal vesicles (3/3 mice, 100%) but not in the ventral (0/3, 0%) prostate (Figure 1E). The 500 µL instillation volume results in the appearance of intense dye staining in the dorsolateral prostate (3/3 mice, 100%), anterior prostate (3/3 mice, 100%), ventral prostate (3/3, 100%), seminal vesicles (2/3 mice, 66.67%), and ureters (2/3 mice, 66.67%) (Figure 1F).

We conclude that urethral instillation fluid volumes ≤50 µL do not consistently distribute to the prostate. Prostate inflammation caused by instilled *E. coli* solution volumes of ≤50 µL likely occurs from bacteria entering prostatic ducts through reflux voiding or through swimming, swarming, or twitching motility. Urethral instillation volumes ≥100 µL directly infiltrate the prostate. A volume of 100 µL distributes specifically to the dorsolateral prostate, while 200 and 500 µL volumes distribute to multiple lobes and the 500 µL also distributes to ureters. We do not recommend instilling ≥500 µL fluid into the urethra unless infection of the ureters and ascending infection of the kidneys is desired.

### The concentration of *E. coli* in instillation fluid determines acute prostatic CFU load

A range of *E. coli* concentrations are delivered transurethrally by research groups to study mouse prostatic inflammation (6, 39). Here we tested if the *E. coli* UTI89 concentration in instillation fluid determines the severity of prostatic infection and we determined the minimum concentration for significant infection of the dorsolateral prostate.

Mice were instilled with 100 µL sterile PBS (OD 0), or PBS containing *E. coli* UTI89 (OD 0.2, 0.4, or 0.8). Mice were euthanized within one minute of instillation and right dorsolateral, left dorsolateral, right ventral, left ventral, right anterior, and left anterior prostate were collected, homogenized, and plated in serial dilution to determine CFUs per mL prostatic tissue homogenate. Two to five mice were instilled per group and six tissue homogenates were prepared from each mouse.

An inoculate of OD 0.8 results in more bacterial CFUs in dorsolateral prostate than an inoculate of OD 0.2 (p = 0.0017, Figure 2A). An inoculate of OD 0.4 results in more CFUs in dorsolateral (p = 0.0170, Figure 2A) and anterior (p = 0.0497, Figure 2C) prostate than an inoculate of OD 0.2. Inoculates above OD 0.4 do not result in more CFUs than an inoculate of OD 0.4 in any lobes (Figure 2). Additionally, there is an overall trend toward an increase in CFUs in the left hemi prostate lobes compared to right hemi prostate lobes (p = 0.0804, Figure 2D).

Our findings suggest a threshold bacterial concentration in the instillation fluid (OD 0.4 for *E. coli* UTI89) is needed to achieve significant prostate infection and further increases in the instillation fluid bacterial concentration (up to OD 0.8) do not increase CFUs in the prostate. The dorsolateral prostate is susceptible to *E. coli* UTI89 colonization and is also the lobe where instillation fluid is most readily distributed.

### Free catch urine culture can be used to predict prostatic inflammation

Prostate inflammation in men presents as acute, chronic, or episodic (45), raising a need to determine molecular and cellular phenotypes during inflammation and after resolution. It is particularly challenging to study the consequences of inflammation after resolution because prostate inflammation from transurethral instillation of *E. coli* is variable in penetrance and severity. We tested whether bacterial load in free catch urine, captured one and seven days after transurethral instillation of *E. coli* UTI89, could be used as a non-invasive biomarker to predict which mice will develop or have prostate inflammation.

Urine was collected and cultured at one day and seven days post-instillation and assessed for dorsal prostate inflammation. Detection of >10,000 CFUs of *E. coli* UTI89 in free catch urine culture at 24 hours (the clinical cutoff for bacterial infection) had a sensitivity (90%), specificity (86%), positive predictive value (90%), and negative predictive value (86%) for dorsal prostate inflammation. Detection of any CFUs of *E. coli* UTI89 in free catch urine culture at seven days had a sensitivity (70%), specificity (57%), positive predictive value (70%), and negative predictive value (57%) for dorsal prostate inflammation.

A free catch urine culture at 24 hours post bacterial instillation appears to be an effective non-invasive biomarker to predict which mice will display dorsal prostate inflammation six days later. We used current clinical recommendations of cutoffs for diagnosing urinary tract infection in veterinary practice, and determining bacterial load in free catch urine provides a robust means for tracking mice with a probable inflammatory event over time, including timepoints after inflammation and colonization have resolved.

### 100 µL of *E. coli* UTI89 (OD 0.80) contains 1-8.2 x10^8^ CFUs of *E. coli*

A generic *E. coli* growth curve is often used to convert OD to CFU concentrations and reported as a single value for the number of *E. coli* colonies delivered in a transurethral bolus. Such a practice fails to account for bacterial strain specific growth curves, individual lab growth conditions, and masks the inherent biologic variability of the model. To determine the range of CFUs for *E. coli* UTI89 culture and transurethral instillation, and to establish biologically significant cutoffs for CFUs in tissue or urine, we measured the concentration of twelve (six per tube) instillation aliquots originating from two separate culture tubes.

A loop of frozen *E. coli* UTI89 was streaked for isolation on a LB KAN agar plate and incubated at 37°C for 24 hours. Streaked plates were stored at 4°C for up to two months. One day prior to instillation a single colony of *E. coli* UTI89 was transferred to 5 mL of antibiotic free LB broth and incubated for 18 hours at 37°C. The optical density was then determined using 1 mL of culture solution. The remaining 4 mL of culture solution was centrifuged at 1157 rcf for 15 minutes at 25°C. Supernatant was removed with gentle suction, taking care not to disturb the *E. coli* UTI89 pellet. The pellet was suspended in sterile PBS. Two culture tubes (Tube 1 and Tube 2) of inoculation solution at an OD of 0.80 were prepared by the same person on the same day and both colonies were taken from a single streaked plate. CFUs per 100 µL sample ranged from 1-8.2 x 10^8^ CFUs. The difference between tubes approached significance (mean of 2.06 x 10^8^ in Tube 1 and 5.33 x 10^8^ in Tube 2, p = 0.0584, Figure 3). Although the two tubes approached significantly different sample CFU counts, they remained within one order of magnitude of each other.

Based on our results we recommend using a cutoff of at least one order of magnitude a difference when comparing CFU counts between groups, as differences less than this may be due to the intrinsic variability of inoculate samples rather than true biologic differences.

### Larger inoculation volumes and *E. coli* concentrations do not change the incidence of bleeding, leaking, or irregular voiding associated with catheter placement

A criticism of delivering *E. coli* to establish prostate inflammation and alter urinary voiding behavior is that it requires catheterization and instillation of fluid. There is a concern that fluid loading the urinary tract will influence voiding by damaging the bladder, prostate, and urethra. Because of this concern, some researchers utilize small volumes of instillation fluid and rely on reflux urine or swimming, swarming, or twitching of bacteria to attain prostate infection. Other researchers use a large instillation volume to directly infuse bacteria into the prostate. We used two methods of bacterial instillation: one low volume (50 µL of sterile PBS containing *E. coli* UTI89 OD 0.35, which we expect does not deliver bacteria directly into the prostate based on outcomes of dye instillation studies in Figure 1), and one high volume (100 µL of sterile PBS containing *E. coli* UTI89 OD 0.7, which we expect directly infects the prostate lobe based on the dye instillation outcomes in Figure 1). Using these two methods, we examined the influence of the inoculation method on the incidence of catheter associated bleeding and leaking, the predicted efficiency of attaining dorsal prostate inflammation using free catch urine culture, and urinary function as measured by void spot assay one, three, five, and seven days post instillation.

We collected free catch urine for culture one- and seven-days post instillation and plated 10 µL of urine on LB KAN+ plates. Presence of greater than 10,000 CFU/mL was considered a “clinical” infection one day post instillation. Presence of any colonies after 24 hours incubation at 37°C was considered positive at seven days post instillation. There is not a significant association between “clinical” infection and instillation method at one day post instillation. 54.55% of mice instilled with 100 µL of sterile PBS containing *E. coli* UTI89 with OD 0.7 are positive at seven days compared to 25.00% in mice instilled with 50 µL of sterile PBS containing *E. coli* UTI89 with OD 0.35 (p = 0.3521). However, there is a significant association between presence of positive culture and the *E. coli* UTI89 infection method at seven days. 81.82% of mice instilled with 100 µL of sterile PBS containing *E. coli* UTI89 (OD 0.7) are positive at seven days compared to 28.57% of mice instilled with 50 µL of sterile PBS containing *E. coli* UTI89 (OD 0.35) (p = 0.0491).

We used a scale of 0-3 to score leaking of instillation fluid from the catheter and urethral bleeding associated with catheterization. For leaking: 0-absent, 1 <10%, 2 <50%, and 3 ≥50% of instilled volume. For bleeding: 0-absent, 1-minimal, but no continuous bleeding, 2-minimal continued bleeding, 3-continued bleed that requires holding pressure. There are no significant differences between the four groups for either leaking or bleeding scores (Figure 4). A larger instillation fluid volume (100 µL) containing *E. coli* UTI89 does not increase catherization associated bleeding and leaking. These findings support use of the higher volume instillation volume of 100 µL with *E. coli* UTI89 OD > 0.40 without concern for increased confounding trauma in C57Bl/6J mice.

Researchers in the field have expressed concerns about using transurethral infections to drive prostate inflammation because it had been thought that fluid loading caused by large volume of instillation solution would by itself elicit changes in voiding and confound studies. We compared void spot assay parameters between mice instilled with a low volume of fluid, a high-volume of fluid, and their respective controls. Detailed void spot assay results are in Table 1-4, 7 and summarized in Table 5. We detected no significant differences between mice instilled with either 50 or 100 µL sterile PBS control groups at any time and conclude that transurethral catheterization and fluid loading with up to 100 µL of sterile PBS does not significantly change adult male mouse urinary function.

**Table 6.** Bladder pressure associated with filling and emptying seven to eight d post urethral instillation with sterile PBS (control) or PBS containing *E. coli* UTI89.

| **Cystometry** | Values are: Average (SD) |  |  |
| --- | --- | --- | --- |
|  | **Sterile PBS**<br>**Control**<br>**(n = 12)** | **PBS w/ *E. coli***<br>**UTI89**<br>**(n = 12)** | **p-value** |
| Void Duration | 0.7465 (0.2880) | 0.7033 (0.2457) | 0.6272 |
| Intervoid Interval | 4.814 (1.792) | 4.866 (1.762) | 0.9254 |
| Baseline Pressure | 2.652 (0.9032) | 2.782 (0.9265) | 0.7261 |
| Normalized Threshold Pressure | 2.677 (1.108) | 3.082 (0.9127) | 0.3276 |
| Normalized Peak Void<br>Pressure | 19.26 (6.043) | 20.53 (4.999) | 0.5717 |
| Non-voiding contractions | 3.417 (2.072) | 3.890 (1.459) | 0.5131 |
| Voided Volume ($\times 10^{-2}$) | 0.06622<br>(0.02635) | 0.06527<br>(0.02294) | 0.8061 |
| Compliance ($\times 10^{-2}$) | 0.02269<br>(0.01012) | 0.01978<br>(0.008291) | 0.2863 |
| Volume Flow Rate ($\times 10^{-3}$) | 0.001715<br>(0.0008992) | 0.001814<br>(0.0008551) | 0.7813 |
| Mass Based Flow Rate | 37.80 (9.447) | 41.18 (11.03) | 0.3134 |
| Efficiency (%) | 105.0 (19.90) | 103.9 (11.35) | 0.3684 |

**Table 7.**
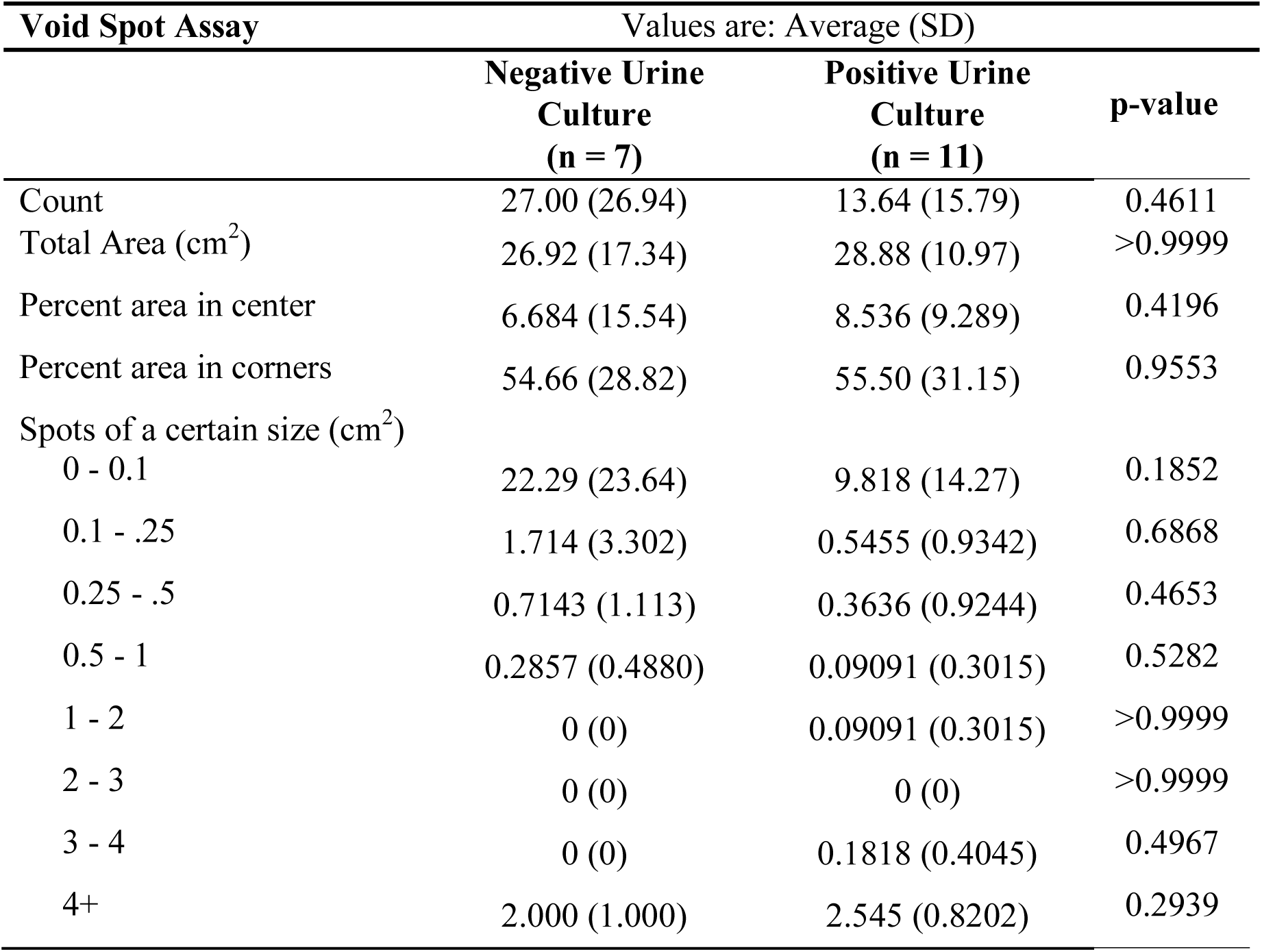
Spontaneous void behavior 7d post urethral instillation positive vs. negative urine culture.

At three, five, and seven days post instillation there is a difference between one of the *E. coli* UTI89 groups and one of the control groups consistent with high volume low frequency voiding in infected mice. Three days post-instillation, mice instilled with 100 µL of sterile PBS containing *E. coli* UTI89 (OD 0.7) void less frequently than mice with 50 µL of sterile PBS (control, p = 0.0050). The difference in voiding frequency derives specifically from small volume void spots (0-0.1 cm^2^, p = 0.0060). Five days post instillation, mice instilled with 50 µL of sterile PBS containing *E. coli* UTI89 (OD 0.35) void in the corners of the cage less frequently than mice instilled with 100 µL of sterile PBS control (p = 0.0211). Seven days post instillation, mice instilled with 100 µL of sterile PBS containing *E. coli* UTI89 (OD 0.7) void less 0.25-0.5 cm^2^ spots than mice instilled with 100 µL of sterile PBS control (p = 0.0250). Also, at seven days, mice instilled with 100 µL of sterile PBS containing *E. coli* UTI89 (OD 0.7) have larger void spots (>4 cm^2^) than mice instilled with either 50 or 100 µL of sterile PBS (p = 0.0018 and 0.0300).

The volume and concentration of *E. coli* UTI89 instilled into the urethra influences spontaneous voiding behaviors seven days post instillation. Mice instilled with 100 µL of sterile PBS containing *E. coli* UTI89 (OD 0.7) void less frequently than mice instilled with 50 µL of sterile PBS containing *E. coli* UTI89 (OD 0.35) (p = 0.0366). The difference in voiding frequency is driven specifically by small volume void spots, specifically spots that are 0-0.1cm^2^ (p = 0.0043) and spots that are 0.25-0.5cm^2^ (p = 0.0136).

Due to the increased percentage of mice with a positive urine culture at seven days in mice instilled with 100 µL of sterile PBS containing *E. coli* UTI89 (OD 0.7) compared to 50 µL of sterile PBS containing *E. coli* UTI89 (OD 0.35), we tested whether the voiding pattern of mice with an *E. coli* UTI89 positive urine culture at seven days post infection differs from that of mice with a negative urine culture at seven days. There were no significant differences between those with a positive urine culture and those with a negative urine culture (Table 7).

We conclude that mice instilled with 100 µL of sterile PBS containing *E. coli* UTI89 (OD 0.7) and mice instilled with 50 µL of sterile PBS containing *E. coli* (OD 0.35) have a similar voiding phenotype one to five days post instillation, showing a largely preserved acute voiding phenotype between the two instillation methods. We hypothesized that voiding differences at seven days were due to an increased percentage of mice with positive urine culture in the group instilled with 100 µL of sterile PBS containing *E. coli* (OD 0.7) but this hypothesis was rejected as there are no differences in voiding phenotype between mice with a positive and negative urine culture at the seven day timepoint. This is particularly interesting because it supports the idea that exposure to a different volume/concentration of *E. coli* UTI89 is sufficient to change duration of voiding phenotypes and is independent of *E. coli* burden. It is possible that any exposure to *E. coli* creates a durable response that persists at least one-week. However, further study is needed on the coordinated response to *E. coli* exposure.

### *E. coli* UTI89 infection results in a low frequency, high volume voiding phenotype

We grouped all control mice and all *E. coli* UTI89 infected mice, to maximize sample size, and conducted a pairwise comparison of spontaneous void spot assays endpoints one day, three days, five days, and seven days post infection and cystometry endpoints seven-eight days post infection. Detailed results are in Table 1-4, 6 and results are summarized in Figure 5. At one day post-instillation there are no significant voiding differences between *E. coli* infected and PBS controls. At three days post instillation, *E. coli* UTI89 infected mice void less frequently than PBS controls (p = 0.0436). At five days post instillation, *E. coli* UTI89 infected mice void a greater volume of urine (p = 0.0301) and void less frequently in the corners of the cage (p = 0.0192) than PBS controls. At seven days post instillation, *E. coli* UTI89 infected mice void a greater volume of urine (p = 0.0044) and void more frequently in large volumes (greater number of spots 4 cm^2^ in area or larger, p = 0.0011) than PBS controls. Abridged VSA results are depicted in Figure 5 and compare *E. coli* UTI89 infected mice to PBS control over the seven-day period. There are no significant differences in cystometric endpoints between *E. coli* UTI89 instilled and PBS control at seven-eight days post instillation.

To test whether transurethral catheterization influences urinary physiology, we compared void spot assay results obtained seven days after sterile PBS instillation to results from historical control mice of the same age and strain that were not catheterized (42). We found no differences between mice given a transurethral catheter and instilled with sterile PBS and mice that were not catheterized. Mice instilled with *E. coli* UTI89 void a significantly greater volume of urine and void more frequently in large volumes than mice that were catheterized and given sterile PBS and mice that were not catheterized (Figure 6). We also compared cystometry results at seven days post sterile PBS instillation to that of age and strain matched historical controls that were not catheterized (42) and detect no significant differences.

**Figure 6.**
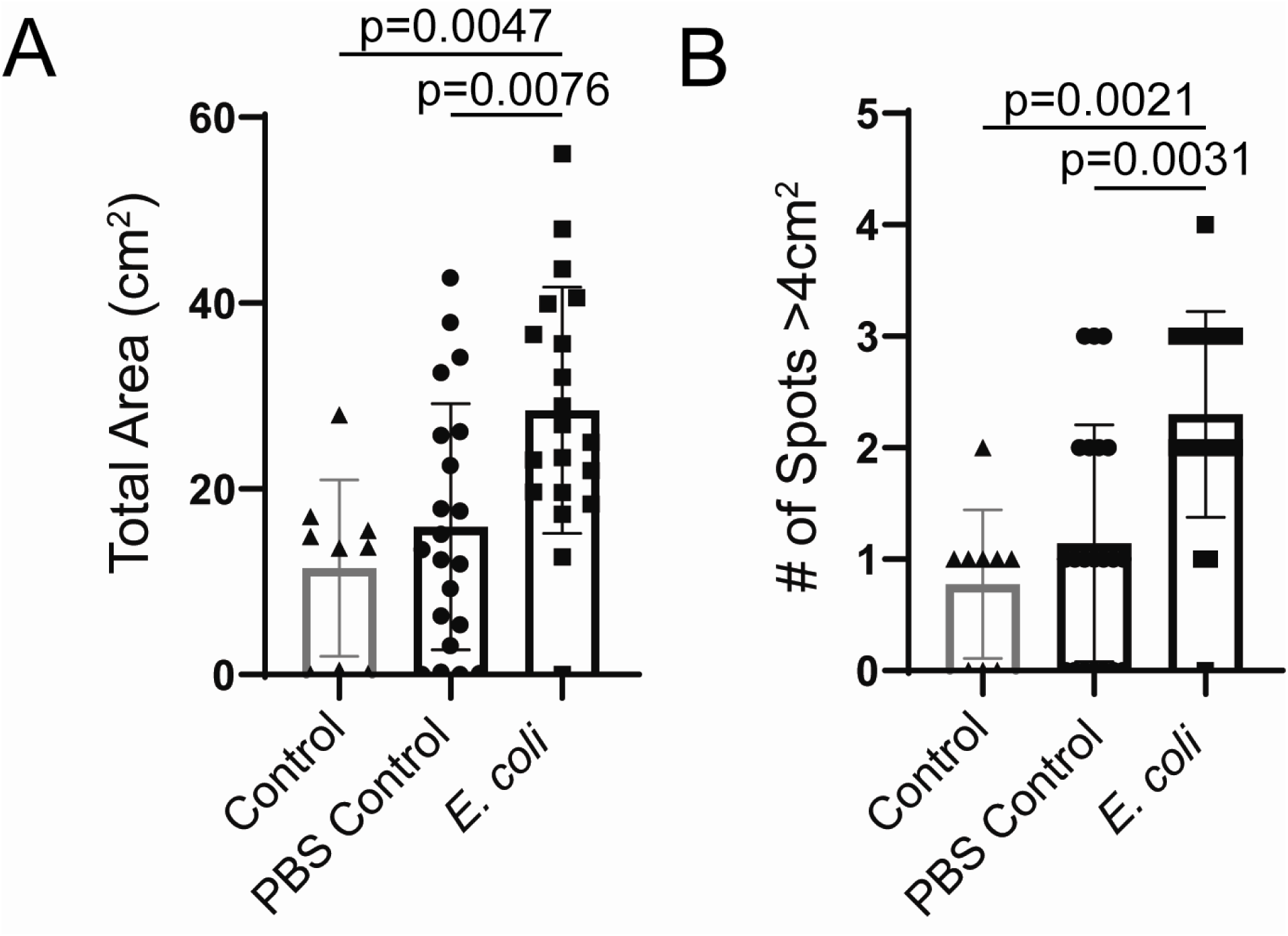
Catheterization alone does not impact voiding. We compared void spot assay results obtained seven days after sterile PBS instillation to results from historical control C57BL/6J male mice of the same age and strain that were not catheterized (42). Graphs are mean ± SD and representative of n = 9-21 per group. A Shapiro-Wilk test was used to test for normality and transformation was applied to normalize data. Bartlett’s test was used to test for homogeneity of variance. When variance was equal, comparisons between groups were made using ordinary one-way ANOVA followed by Tukey’s multiple comparisons test. If data could not be normalized through transformation a Kruskal-Wallis test was applied. A p < 0.05 was considered statistically significant.

It is interesting that *E. coli* UTI89 infected mice display a high volume, low void frequency phenotype. This voiding phenotype suggests increased water intake based on the high-volume phenotype in *E. coli* UTI89 infected mice but does not explain the low void frequency phenotype. The high-volume void is also consistent with a physiologic response of high-volume voids to flush the urinary tract. The low void frequency may suggest a reluctancy to urinate.

When using cystometry to evaluate voiding behavior, no differences were observed between infected and uninfected mice. One possible explanation is that the VSA is more sensitive than cystometry in detecting bacterial-induced changes to urinary voiding. Another possibility is that the bacterial change in voiding phenotype is a voluntary behavioral response that is eliminated during anesthesia. Butler et al. determined that social stress could cause long lasting voiding dysfunction in mice, specifically less frequent and larger volume voids (8). *E. coli* exposure and instillation of PBS may trigger a stress pattern of voiding like what is manifested during aggressive behavior.

The voiding response to bacterial infection that we observed differs from the response reported in previous work. Specifically, previous studies identified a low volume, high void frequency phenotype with acute *E. coli* 1677 infection (25). The results described here may differ from previous studies because we used a different strain of mice, a different strain of *E. coli*, a different installation volume and a different way to quantify *E. coli* and void spot assay papers. C3H/HeOuJ mice used in a previous prostate infection study harbor an toll receptor variant (lps^d^) that renders them hyporesponsive to endotoxin (18, 46) and makes them susceptible to chronic infection (34). Prostatic *E. coli* infections in C3H/HeOuJ mice are persistent and become lymphocytic over time and this may lead to a different voiding phenotype than prostate infections in C57Bl/6J mice. Some research groups omit small spots from analysis attributing them to footprints, though our previous study supports the notion that impact of mouse tracking through deposited urine spots is negligible (51). We used Void Whizzard software (51) and followed recommendations for the reporting of void spot assay data (19) to improve the rigor and reproducibility of our void spot assay findings. Further studies are needed to parse out driving factors of this differed voiding phenotype.

### Transurethral instillation of *E. coli* UTI89 increases dorsal prostate lobe stromal cell density and collagen content

Histological inflammation is confined to the dorsal prostate, so collagen density was quantified exclusively in this lobe. Histological inflammation is extremely variable between mice with some *E. coli* UTI89 infected mice displaying no inflammation and others with expansive leukocyte recruitment and infiltration (Figure 7C). Picrosirius red staining was conducted to assess dorsal prostate collagen content. *E. coli* UTI89 infected mice have significantly denser collagen (p = 0.0221) than sterile PBS control mice (Figure 7A). In addition, there is a strong direct correlation (p < 0.0001) between dorsal prostate stromal cell number and collagen density (Figure 7B).

**Figure 7.**
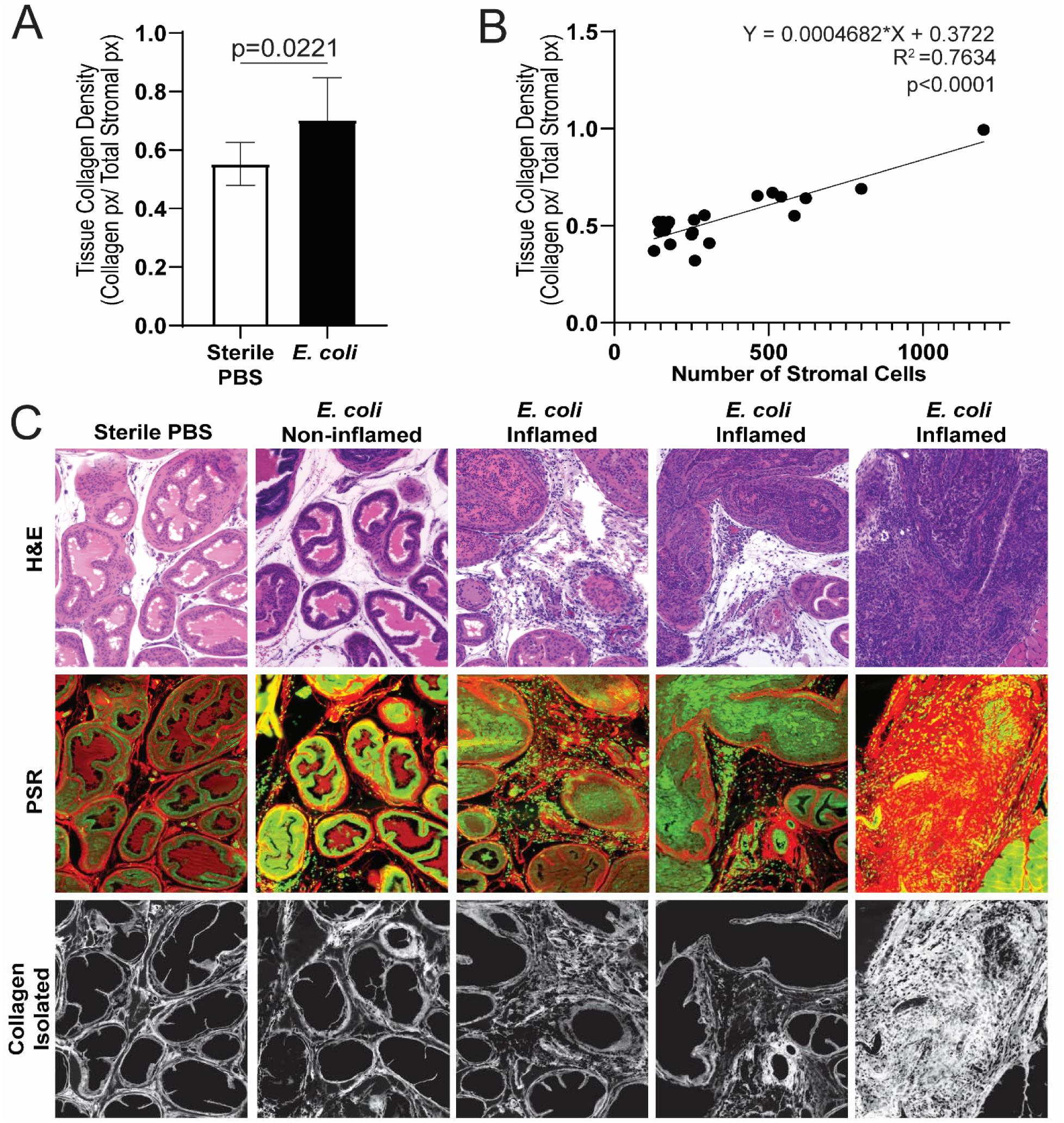
Transurethral instillation of *E. coli* UTI89 into C57BL/6J male mice results in increased dorsal prostate lobe stromal cell density and collagen accumulation. Histological inflammation was confined to the dorsal prostate and collagen density was quantified exclusively in this lobe. (A) Picrosirius red (PSR) staining was conducted to assess dorsal prostate collagen content. Graph is mean ± SD, and representative of 6-7 mice per group. A Mann Whitney test was used to compare groups. (B) Linear regression analysis was performed to compare dorsal prostate stromal cell number and collagen density. Results are representative of assessment of 21 regions of interest. (C) Histological inflammation was variable between mice with some *E. coli* UTI89 infected displaying no inflammation and others with expansive leukocyte recruitment and collagen accumulation.

**Figure 8.**
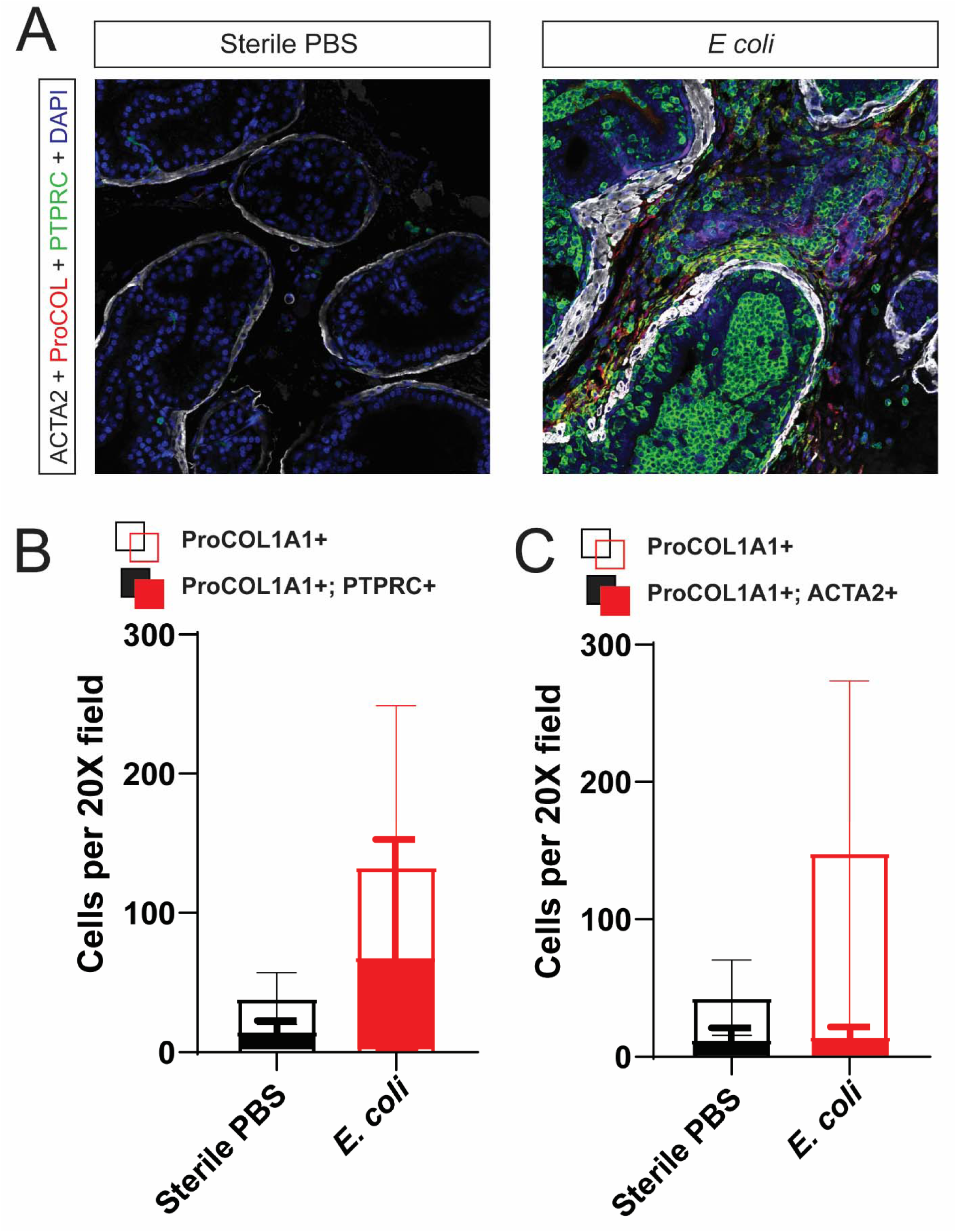
Transurethral instillation of UTI89 increases the density of ProCOL1A1+ cells in the C57BL/6J male mouse prostate. Transurethral instillation of UTI89 increases the density of PTPRC+ ProCOL1A1+ cells and decreases the percentage of ACTA2+ ProCol1A1+ cells in the prostate C57BL/6J male mouse prostate. (A) Immunohistochemical staining for smooth muscle actin (ACTA2), procollagen 1A1 (ProCol1A1), protein tyrosine phosphatases, receptor type C (PTPRC, also known as CD45) was performed on dorsal prostate section from *E. coli* UTI89 infected mice and PBS controls. (B and C) Infection increases the density of total collagen producing cells (ProCol1A1+) compared to the sterile PBS control (p = 0.0369). (B) Infection also increases the density of collagen producing bone-marrow derived cells (p = 0.0420). However, the proportion of collagen producing immune cells (dual PTPRC+; ProCOL1A1+/ Total ProCOL1A1+) is similar between groups (p = 0.1531). (C) Infection does not alter the density of ProCOL1A1+;ACTA2+ cells (p = 0.7233). However, the proportion of collagen producing ACTA2+ cells (dual ACTA2+; ProCOL1A1+/ Total ProCOL1A1+) is significantly lower compared to control (p = 0.0326). Graphs are mean ± SD and representative of six to seven mice per group. A Shapiro-Wilk test was used to test for normality and transformation was applied to normalize data. Bartlett’s test was used to test for homogeneity of variance. When variance was equal, comparisons between groups were made using ordinary one-way ANOVA followed by Tukey’s multiple comparisons test. If data could not be normalized through transformation a Kruskal-Wallis test was applied. A p < 0.05 was considered statistically significant.

Our results support previous findings that transurethral instillation favors infiltration of the dorsal prostate lobe. Bacterial load drives immune cell recruitment and the extent of the immune cell recruitment drives collagen production and accumulation. *E. coli* UTI89 infected mice may exhibit variable responsiveness to inflammation for a variety of reasons. When mice are instilled with *E. coli* UTI89, there is potential variation in the instillation technique from mouse to mouse, and which may vary in the amount of *E. coli* UTI89 in the prostate. In addition, each mouse may have a different biologic response to the *E. coli* UTI89 infection, further varying the immune response. Despite the variability in response, collagen density increases when inflammation is present.

### Transurethral instillation of *E. coli* UTI89 increases the number of ProCOL1A1+ cells. Transurethral instillation of *E. coli* UTI89 increases number of PTPRC+ ProCOL1A1+ cells and decreases the percentage of ACTA2+ ProCol1A1+ cells

E. coli *UTI89 infection increases* density of total collagen producing cells (ProCol1A1+) compared to the sterile PBS control (p = 0.0369). Infection also increases the density of collagen producing bone-marrow derived cells (p = 0.0420). However, the proportion of collagen producing bone marrow derived cells (dual PTPRC+; ProCOL1A1+/ Total ProCOL1A1+) is similar between groups (p = 0.1531). Infection results in a similar density of collagen producing ACTA2+ cells compared to sterile saline-infused controls (p = 0.7233). Furthermore, the proportion of collagen producing ACTA2+ cells (dual ACTA2+; ProCOL1A1+/ Total ProCOL1A1+) is significantly lower in infected mice compared to controls (p = 0.0326).

The fact that *E. coli* UTI89 significantly increases the density of total collagen producing cells indicates that infection triggers a fibrotic response in the dorsal prostate. The significant increase in collagen producing bone-marrow derived cells suggests, but does not confirm, that fibrocytes are recruited to the prostate along with other inflammatory cells and contribute nearly 50% of the collagen producing cells in *E. coli* UTI89 infected prostates. Future lineage analysis could provide definitive evidence of the identity of these collagen 1A1 producing cells. The significantly lower proportion of collagen producing ACTA2+ cells (dual ACTA2+; ProCOL1A1+/ Total ProCOL1A1+) in the *E. coli* UTI89 infected group provides evidence that myofibroblasts are not a major contributor to excess collagen production following *E. coli* UTI89 infection, a contrast to the cellular origins of collagen production in other organs. Future work is needed to determine if this is a timing or insult specific response in the prostate or if the prostate has a truly unique collagen production process.

## CONCLUSIONS

We examined the use of transurethral instillations of UTI89 *E. coli* to induce prostate infection and inflammation in mice and thereby model prostate inflammation in men. We optimized instillation volume, instillation *E. coli* UTI89 concentration and identified a non-invasive biomarker for prostatic inflammation at one week thereby establishing a predictable and reproducible model. We determined that: volume of instillations positively correlates with the extent of inflammation distribution, an observed threshold of OD 0.4 is needed to achieve significant prostate infection, free catch urine culture assays are an effective means to predict if the mouse experience prostatic inflammation, mice that are infected demonstrate a low frequency, high voiding phenotype and transurethral instillations increase collagen density in the prostate, which is associated with an increased total number of ProCol1A1+ cells.

## ACKNOWLEDGEMENTS

The authors would like to thank Petra Popovics and Brandon Scharpf for their engaging discussions and helpful critics during manuscript preparation. Funded by National Institutes of Health grants: U54 DK104310, Summer Program In Undergraduate Urologic Research (U54 DK104310S1), R01ES001332, R01DK099328, F31ES028594, TL1TR002375, F30DK122686 and University of Wisconsin-Madison, School of Veterinary Medicine. THS and JMB are supported by George M. O’Brien Urology Research Center grant 2U54DK104309-06 (to JMB). The content is solely the responsibility of the authors and does not necessarily represent the official views of the National Institutes of Health.

